# Residue-level mapping of crowding effects on protein phase separation

**DOI:** 10.1101/2025.10.02.680141

**Authors:** Wei Chen, Jacob M. Shaffer, Christine D. Keating, Scott A. Showalter

## Abstract

Proteins can undergo liquid-liquid phase separation to form condensates within the crowded cellular environment. *In vitro*, polymer-based crowding agents are widely used to mimic this effect, but how their chemical environments influence phase separation compared to native cytosol remains unclear. Here, we use NMR to probe the chemical influences on the intrinsically disordered region of RNA polymerase II under different crowding conditions, including polymer-based crowders, protein-based crowders, and reconstituted *E. coli* cytosol. We find a general trend of enhanced protein self-interactions across all conditions, but also distinct chemical environments that depend on crowder identity, reflecting changes in preferential interactions. Given the widespread use of polymer crowders in phase separation studies, our results establish a strategy to dissect the microscopic role of crowders and lay the foundation for designing more physiologically relevant *in vitro* crowding models of protein phase separation. More broadly, this framework enables systematic probing of residue-level environmental influences in complex settings including within the cellular milieu.

---

Biomolecular phase separation has emerged as a leading explanation for cellular compartmentalization through the segregation of macromolecules into liquid-like condensates^1^. Phase separation is driven by multivalent intermolecular interactions that are often mediated by intrinsically disordered regions (IDRs) of proteins^2^. *In vitro* studies of phase separation typically focus on purified proteins, and polymers such as polyethylene glycol (PEG), Ficoll, and dextran are frequently added to enhance phase separation. While sequence and compositional determinants of protein phase separation have been extensively studied^3^, how crowding influences the phase behavior of proteins is less well understood. Crowding agents are commonly assumed to be inert and simply occupy space to mimic macromolecular crowding in the cell. However, they can also engage in preferential interactions^4^ and even co-condense with a variety of biomolecules^5^. In addition, a large number of reported phase-separating proteins lack data in the absence of crowding agents^6^. Despite their widespread use, the ability of crowders to recapitulate the effects of cellular crowding on protein phase separation, specifically how they shape the chemical environment around proteins, remains largely unexplored.

To address this gap, we investigated the phase separation behavior and the chemical environments sensed by the C-terminal domain (CTD) of the RNA polymerase II (Pol II) across different crowding conditions, formed by crowding agents and reconstituted *E. coli* cytosol. The intrinsically disordered CTD has been attributed a significant role in the formation of dynamic Pol II clusters in cells via phase separation. It consists of tandem repeats of the amino acid sequence YSPTSPS (*Figure 1A*), whose repetitive and low-complexity nature enables multivalent interactions for phase separation, while also making the CTD a simple model for probing crowder influences. Phase separation of the CTD has be demonstrated for both the yeast CTD (26 repeats) and human CTD (52 repeats)^7, 8^, with the number of repeats correlating with phase separation tendency *in vitro* and *in vivo*. Polymer crowders such as dextran have been widely used to promote CTD phase separation *in vitro*^7, 8^. To dissect how crowding mediates CTD phase separation, we systematically compared the effects of three categories of crowders— polymers (PEG, Ficoll, and dextran), proteins (bovine serum albumin (BSA) and lysozyme), and reconstituted *E. coli* cytosol, and mapped the microscopic environment they create.

**Figure 1.**
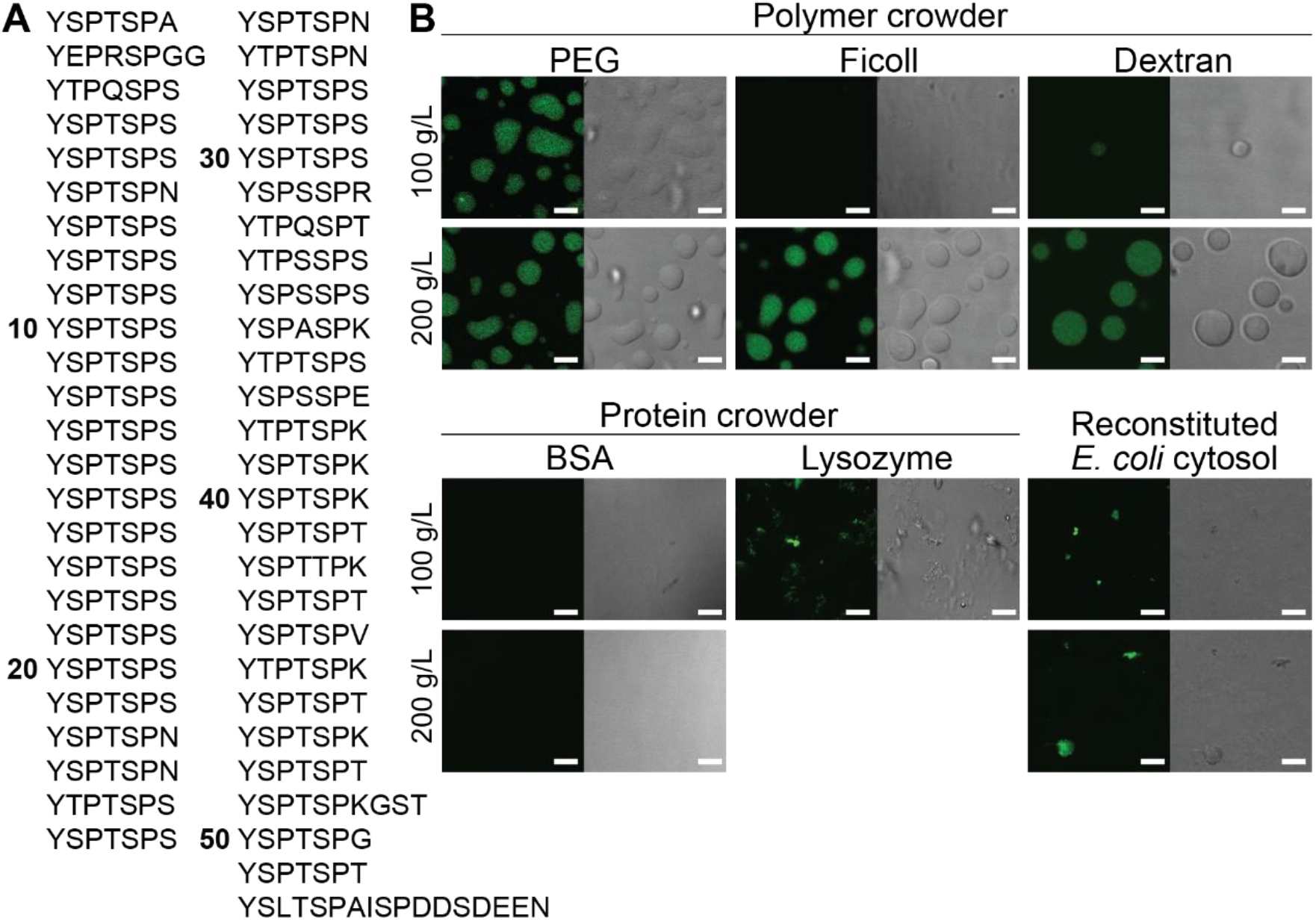
Phase separation behaviors of the human CTD (hCTD) at various crowding conditions. (A) The sequence of hCTD contains 52 tandem heptad repeats with the consensus sequence of YSPTSPS. (B) Confocal fluorescence and differential interference contrast (DIC) imaging showing phase separation of 25 μM MBP-hCTD, fluorescently labeled with Alexa Fluor 488, depends on the identity and concentration of crowders. While polymers promoted CTD droplet formation, proteins and reconstituted *E. coli* cytosol did not. Lysozyme and *E. coli* cytosol led to the formation of irregular CTD aggregates. Scale bar: 5 μm. Experiments were carried out in 50 mM HEPES pH 7.5 and 150 mM NaCl. Fluorescence images were false colored with adjusted brightness/contrast for clarity.

We first compared the phase behavior of the human CTD (hCTD) across different crowding conditions. Among three commonly used polymer crowders, PEG (MW 8 kD) is linear, dextran (MW 9 to 11 kD) is a slightly branched (∼5%), and Ficoll (MW 70 kD) is highly branched. The CTD phase separated into liquid droplets in the presence of all three polymers at a crowder concentration of 200 g/L (*Figure 1B*). PEG was the most effective driver, producing abundance droplets even at 100 g/L, whereas Ficoll had no effect and dextran yielded only rare small droplets. In contrast to polymers, globular proteins are more rigid and can provide nonspecific protein-protein interactions that more closely resemble those in cells. We therefore examined two protein-based crowders—BSA (MW 66 kD, pI 5) and lysozyme (MW 14 kD, pI 9), which are oppositely charged at pH 7.5. Under these conditions, BSA did not promote CTD phase separation, while lysozyme induced irregular aggregate formation. To more closely mimic cellular composition, we reconstituted *E. coli* cytosol at 100 and 200 g dry weight/L from lyophilized lysate, reflecting physiological macromolecular concentrations (100 to 400 g/L). The reconstituted cytosol contained soluble cytoplasmic proteins, nucleic acids, and ions, but excluded cell walls, membrane fragments, and insoluble proteins. Similar to lysozyme, reconstituted cytosol led to CTD aggregation, though on a smaller scale with fewer aggregates. Taken together, these results show that even at the same macromolecular concentration, the identity of the crowder determines CTD phase behavior. Notably, only polymer crowders supported liquid condensate formation under our conditions.

To dissect the molecular basis of crowder-mediated phase separation, we used NMR spectroscopy to map the chemical environments of the CTD at residue-level resolution. NMR has been widely applied to study how crowding influences protein stability^9^, folding^10^, and association^11^, but its potential to elucidate residue-level crowding effects on phase separation remains unrealized. Here, we focused on a shorter CTD construct containing 12 YSPTSPS repeats (CTD12), chosen for its high expression yield and ability to undergo phase separation (520 μM with 300 g/L PEG). The CTD is rich in prolines (∼30%), which are prevalent in other IDRs^12^ and have been implicated in CTD phase separation through contacts with tyrosines and *cis-trans* isomerization^8^. However, prolines are traditionally challenging for protein NMR because they lack amide hydrogens and thus invisible in ^1^H-^15^N-HSQC spectra (*Figure 2A*). To overcome this limitation, we carried out ^13^C direct-detect ^13^C, ^15^N-CON experiments (*Figure 2B*), which allowed a direct visualization of prolines with improved spectral dispersion. We observed both P_3_ and P_6_ in *trans* conformation and also *cis*-P_3_ (10% of P_3_ population). We focused on the internal repeats of CTD12, rather than the terminal ones, as they better represent the full-length CTD. In the CON spectrum, residues with the same numbering in these Y_1_S_2_P_3_T_4_S_5_P_6_S_7_ repeats (excluding termini) clustered as a single peak, reflecting their averaged chemical environment.

**Figure 2.**
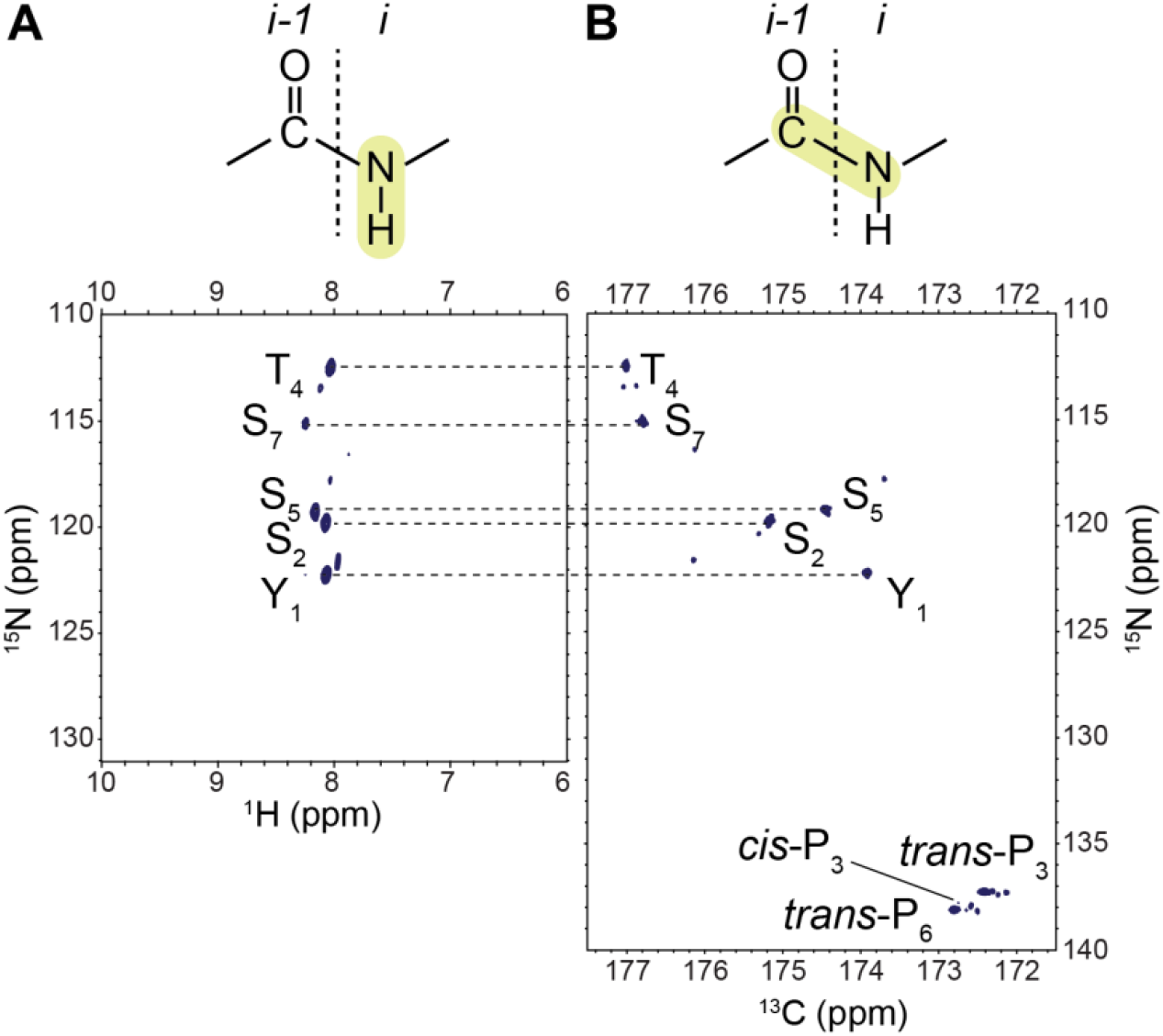
NMR spectra of CTD12 obtained using different detection probes. (A) Proton-detected ^1^H, ^15^N-HSQC with a narrow dispersion in the ^1^H dimension and a lack of proline detection. (B) Carbon-detected ^13^C, ^15^N-CON with a wider dispersion in the ^13^C dimension and direct visualization of proline residues. Experiments were carried out using 425 μM CTD12 in 50 mM HEPES pH 7.5, 150 mM NaCl, 1 mM DSS, and 10% D_2_O.

We first performed ^13^C, ^15^N-CON experiments for CTD12 at two concentrations (425 μM and 1 mM) without crowders and analyzed weighted-average chemical shift perturbations (CSPs) across the ^13^C and ^15^N dimensions. At 1 mM, Y_1_, S_2_, and S_5_ showed the largest concentration-dependent CSPs (*Figure 3A, D*), reflecting CTD self-interactions and consistent with previous studies implicating tyrosines in CTD phase separation^8^ and in the stickers-and-spacers model^13^. We then examined CSPs for 425 μM CTD12 in the presence of 100 mg/mL PEG, dextran, BSA, lysozyme, and reconstituted *E. coli* cytosol. Because crowders are NMR silent, CSPs directly report the chemical environments sensed by the CTD. Crowders produced distinct residue-level CSP magnitudes and patterns (*Figure 3A*), with PEG inducing the largest CSPs (*Figure 3A, C*), consistent with its strong ability to drive CTD phase separation. Notably, *cis*-P_3_ (for which only ^15^N CSP were available) displayed the largest CSP among all residues despite its small population (10% for 425 μM, 9% for 1 mM, 8% for BSA, 4% for PEG, and undetectable for lysozyme and *reconstituted* E. coli cytosol) (Figure *3B, C*). This indicates that *cis*-P_3_ is highly sensitive to concentration and environmental changes and are consistent with previous finding that proline-tyrosine contacts and proline cis-trans isomerization are important for CTD phase separation. Overall, these observations indicate that different crowders create distinct microscopic chemical environments around the CTD.

**Figure 3.**
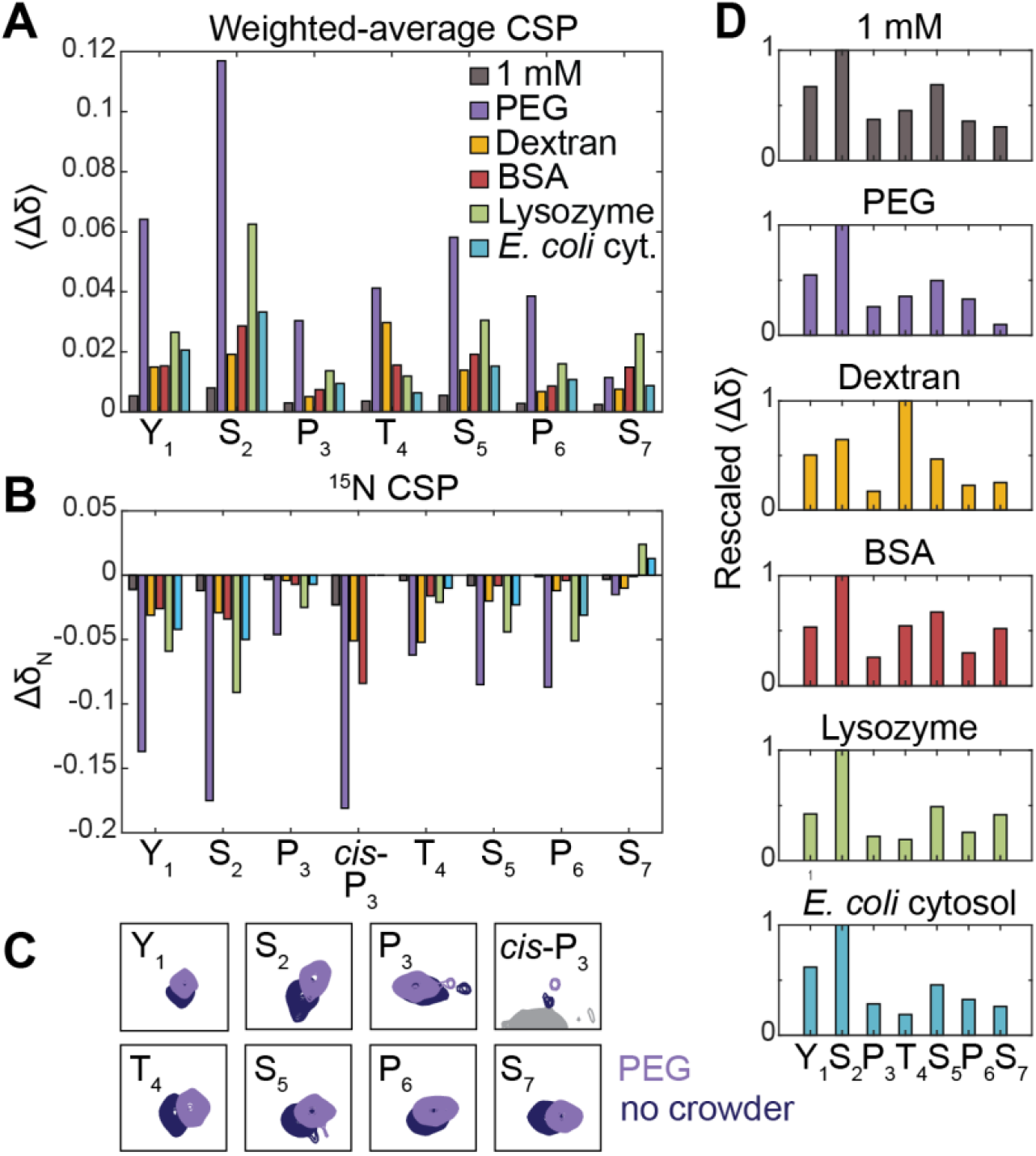
Chemical shift perturbations (CSPs) of CTD12 induced by different crowders. (A) The weighted average of CSPs, ⟨Δδ⟩, for each residue in the repeat unit YSPTSPS reveals that different residues experience distinct chemical environments depending on the identity of the crowder. A higher protein concentration (1 mM, as compared to 425 μM) also caused distinct CSPs for different residues, at a smaller scale. (C) ^15^N CSPs shows large concentration and crowding effects on *cis*-P3. (C) Visualization of peak shifting in the ^13^C, ^15^N-CON spectrum caused by PEG. (D) Rescaled weighted average of CSPs reveals distinct crowder-dependent profiles.

To dissect the contributions to crowder-dependent CSPs, we considered two non-mutually exclusive mechanisms: (1) enhanced CTD self-interactions as a result of crowder volume exclusion, and (2) changes in preferential chemical interactions between the CTD, crowder, and water. To differentiate between these mechanisms, we rescaled the weighted-average CSP profile for each condition by setting the largest value to one (*Figure 3C*) and compared across conditions. If a crowder is inert and acts only through volume exclusion, leading to a higher effective CTD concentration, we would expect its rescaled CSP profile to resemble that of the 1 mM CTD without crowders. Indeed, all crowding conditions, except dextran, exhibited the largest CSP at S_2_, followed by Y_1_ or S_5_, similar to the 1 mM condition, suggesting that crowding increased the effective CTD concentration and enhanced CTD self-interactions. On the other hand, each crowder also differed in their CSP profile—while PEG had less effect at S_7_, lysozyme had more effect at S_7_ and less at T_4_, and *E. coli* lysate had less effect at T_4_, when compared to the 1 mM CTD concentration condition (*Figure 3C*). These differences suggest that, in addition to volume exclusion, there were distinct weak interactions, either associative or repulsive, between the CTD and crowders that shaped the chemical environment experienced by the protein. Dextran deviated the most in terms of its CSP profile compared to that of 1 mM CTD. It exhibited largest CSP at T_4_, unlike in other conditions where it peaked at S_2_, suggesting that chemical interactions played a large role in dextran’s crowding effect on the CTD.

We showed that crowder identity shaped the residue-level chemical environment experienced by an IDR and had drastic effects on protein phase separation. Using NMR, we differentiated crowding-enhanced protein self-interactions and changes in protein/crowder/solvent preferential interactions. Because polymers often poorly recapitulate cellular crowding^14^, more complex mixtures and cell lysates are increasingly used^15^. Given the common use of polymer crowders in phase separation studies, our results provide a foundation for designing more physiologically relevant *in vitro* models by allowing direct comparison to cell-like environments such as nuclear or cytoplasmic extracts, and ultimately to cellular environments via in-cell NMR^16^. More broadly, this work presents a proof of concept for probing residue-level environmental influence, including those in the cellular milieu where physiological crowding and native interactions are present, to systematically decode the molecular grammar for protein phase separation.

## ACKNOWLEDGEMENT

We thank the CSL Behring Fermentation Facility at Penn State for helping prepare the lyophilized *E. coli* lysate, Sarah Chapman for discussion and helping with confocal imaging in the early exploration stage of the project, and Chi-Yun Lin and Lola Sibaud for access to size exclusion chromatography and their assistance. This work was supported by funding from the U.S. National Science Foundation (MCB-1932730 and MCB-2428716 to S.A.S., and MCB-2317529 to C.D.K). W.C. acknowledges support from the Eberly Postdoctoral Research Fellowship. C.D.K. acknowledges support from the Shapiro Professorship in Chemistry.

---

For Table of Contents Only

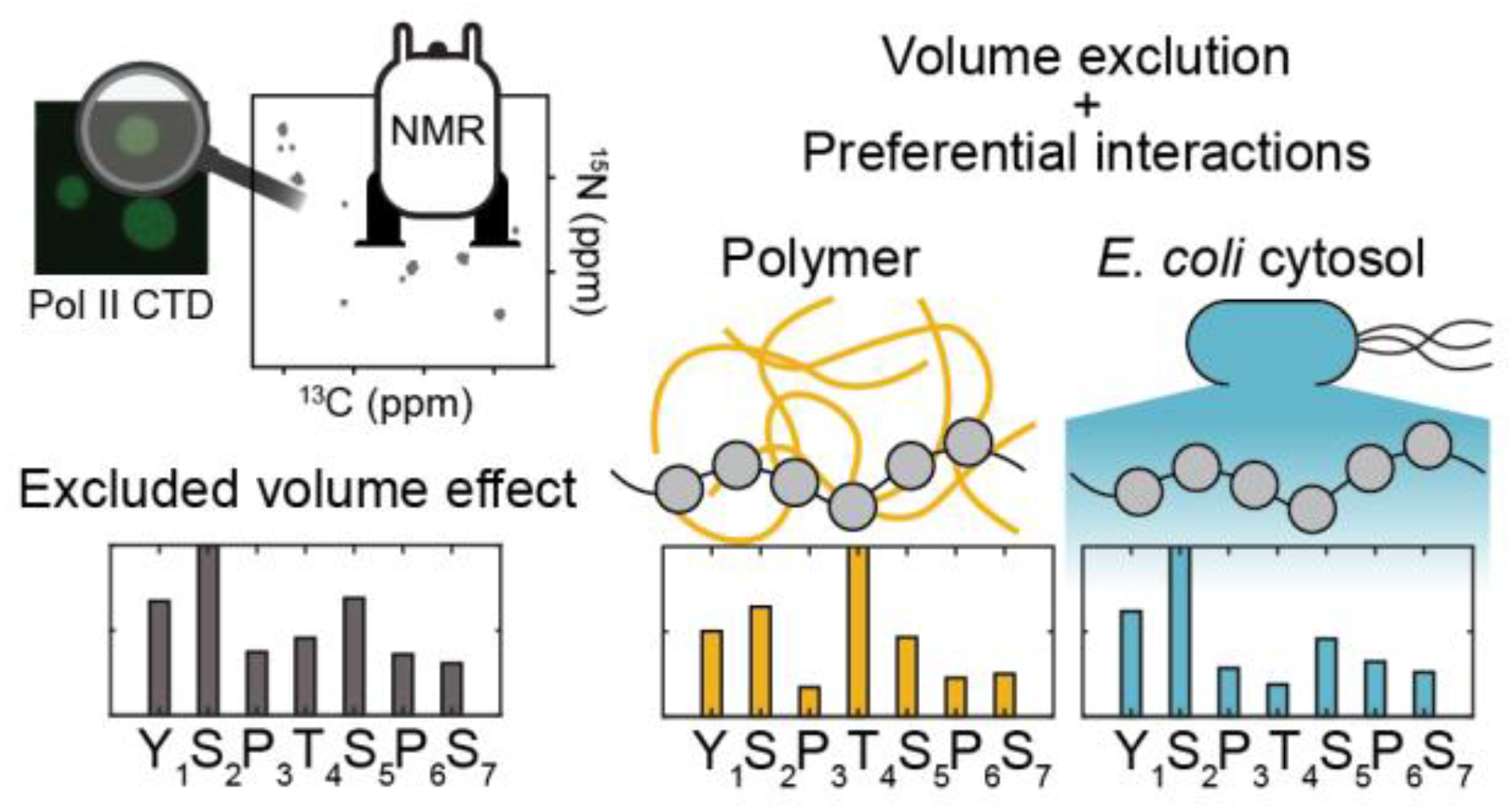

## REFERENCES

(1) Banani, S. F.; Lee, H. O.; Hyman, A. A.; Rosen, M. K. Biomolecular condensates: organizers of cellular biochemistry. Nat Rev Mol Cell Biol 2017, 18 (5), 285–298. DOI: 10.1038/nrm.2017.7.

(2) Chen, W.; Fraser, O. A.; George, C.; Showalter, S. A. From molecular descriptions to cellular functions of intrinsically disordered protein regions. Biophys Rev (Melville) 2024, 5 (4), 041306. DOI: 10.1063/5.0225900.

(3) Vendruscolo, M.; Fuxreiter, M. Towards sequence-based principles for protein phase separation predictions. Curr Opin Chem Biol 2023, 75, 102317. DOI: 10.1016/j.cbpa.2023.102317.

(4) Lee, J. C.; Lee, L. L. Preferential solvent interactions between proteins and polyethylene glycols. J Biol Chem 1981, 256 (2), 625–631. From NLM Medline. Wang, Y.; Annunziata, O. Comparison between protein-polyethylene glycol (PEG) interactions and the effect of PEG on protein-protein interactions using the liquid-liquid phase transition. J Phys Chem B 2007, 111 (5), 1222–1230. DOI: 10.1021/jp065608u.

(5) Marianelli, A. M.; Miller, B. M.; Keating, C. D. Impact of macromolecular crowding on RNA/spermine complex coacervation and oligonucleotide compartmentalization. Soft Matter 2018, 14 (3), 368–378. DOI: 10.1039/c7sm02146a. Qian, D. Y.; Welsh, T. J.; Erkamp, N. A.; Qamar, S.; Nixon-Abell, J.; Krainer, G.; St George-Hyslop, P.; Michaels, T. C. T.; Knowles, T. P. J. Tie-Line Analysis Reveals Interactions Driving Heteromolecular Condensate Formation. Phys Rev X 2022, 12 (4). DOI: ARTN 041038 10.1103/PhysRevX.12.041038. André, A. A. M.; Yewdall, N. A.; Spruijt, E. Crowding-induced phase separation and gelling by co-condensation of PEG in NPM1-rRNA condensates. Biophys J 2023, 122 (2), 397–407. DOI: 10.1016/j.bpj.2022.12.001.

(6) Andre, A. A. M.; Spruijt, E. Liquid-Liquid Phase Separation in Crowded Environments. Int J Mol Sci 2020, 21 (16). DOI: 10.3390/ijms21165908.

(7) Boehning, M.; Dugast-Darzacq, C.; Rankovic, M.; Hansen, A. S.; Yu, T.; Marie-Nelly, H.; McSwiggen, D. T.; Kokic, G.; Dailey, G. M.; Cramer, P.; et al. RNA polymerase II clustering through carboxy-terminal domain phase separation. Nat Struct Mol Biol 2018, 25 (9), 833–840. DOI: 10.1038/s41594-018-0112-y. Zhang, Q.; Kim, W.; Panina, S. B.; Mayfield, J. E.; Portz, B.; Zhang, Y. J. Variation of C-terminal domain governs RNA polymerase II genomic locations and alternative splicing in eukaryotic transcription. Nat Commun 2024, 15 (1), 7985. DOI: 10.1038/s41467-024-52391-6.

(8) Flores-Solis, D.; Lushpinskaia, I. P.; Polyansky, A. A.; Changiarath, A.; Boehning, M.; Mirkovic, M.; Walshe, J.; Pietrek, L. M.; Cramer, P.; Stelzl, L. S.; et al. Driving forces behind phase separation of the carboxy-terminal domain of RNA polymerase II. Nat Commun 2023, 14 (1), 5979. DOI: 10.1038/s41467-023-41633-8. Linhartova, K.; Falginella, F. L.; Matl, M.; Sebesta, M.; Vacha, R.; Stefl, R. Sequence and structural determinants of RNAPII CTD phase-separation and phosphorylation by CDK7. Nat Commun 2024, 15 (1), 9163. DOI: 10.1038/s41467-024-53305-2.

(9) Wang, Y.; Sarkar, M.; Smith, A. E.; Krois, A. S.; Pielak, G. J. Macromolecular crowding and protein stability. J Am Chem Soc 2012, 134 (40), 16614–16618. DOI: 10.1021/ja305300m.

(10) Tokuriki, N.; Kinjo, M.; Negi, S.; Hoshino, M.; Goto, Y.; Urabe, I.; Yomo, T. Protein folding by the effects of macromolecular crowding. Protein Sci 2004, 13 (1), 125–133. DOI: 10.1110/ps.03288104.

(11) Guseman, A. J.; Pielak, G. J. Cosolute and Crowding Effects on a Side-By-Side Protein Dimer. Biochemistry 2017, 56 (7), 971–976. DOI: 10.1021/acs.biochem.6b01251.

(12) Uversky, V. N. The alphabet of intrinsic disorder: II. Various roles of glutamic acid in ordered and intrinsically disordered proteins. Intrinsically Disord Proteins 2013, 1 (1), e24684. DOI: 10.4161/idp.24684.

(13) Martin, E. W.; Holehouse, A. S.; Peran, I.; Farag, M.; Incicco, J. J.; Bremer, A.; Grace, C. R.; Soranno, A.; Pappu, R. V.; Mittag, T. Valence and patterning of aromatic residues determine the phase behavior of prion-like domains. Science 2020, 367 (6478), 694–699. DOI: 10.1126/science.aaw8653.

(14) Tyrrell, J.; Weeks, K. M.; Pielak, G. J. Challenge of mimicking the influences of the cellular environment on RNA structure by PEG-induced macromolecular crowding. Biochemistry 2015, 54 (42), 6447–6453. DOI: 10.1021/acs.biochem.5b00767.

(15) Davis, C. M.; Deutsch, J.; Gruebele, M. An in vitro mimic of in-cell solvation for protein folding studies. Protein Sci 2020, 29 (4), 1060–1068. DOI: 10.1002/pro.3833. Sarkar, M.; Smith, E.; Pielak, G. J. Impact of reconstituted cytosol on protein stability. Proc Natl Acad Sci U S A 2013, 110 (48), 19342–19347. DOI: 10.1073/pnas.1312678110.

(16) Theillet, F. X.; Luchinat, E. In-cell NMR: Why and how? Prog Nucl Magn Reson Spectrosc 2022, 132-133, 1–112. DOI: 10.1016/j.pnmrs.2022.04.002.

