## Supplementary Information for "Residue-level mapping of crowding effects on protein phase separation"

### MATERIALS AND METHODS

#### Protein expression and purification

*MBP-tagged Human CTD (MBP-hCTD).* A pMalX plasmid containing the DNA sequence encoding residues 1592-1970 of the human Rpb1 and a C-terminal 6xHis tag was used. BL21 (DE3) cells containing the hCTD plasmid were grown LB media with 100 μg/mL ampicillin at 37 °C and 200 rpm. At an $OD_{600}$ of 0.7-0.8, expression was induced with 0.3 mM IPTG at 11 °C and 200 rpm for 16 hours. Cells were pelleted by centrifugation at 5000x g and stored at -80 °C. Cell pellets of MBP-hCTD were resuspended in the lysis buffer (50 mM Tris pH 7.5, 500 mM NaCl, 20 mM imidazole) supplemented with 1x EDTA-free protease inhibitor cocktail (Millipore), 1 mM PMSF, and 10 units/L growth RNAse-free DNAse (NEB). Following sonication on ice, the cell lysate was centrifuged at 18000x g for 45 minutes. The supernatant was passed through Ni-NTA resins (Thermo Fisher Scientific) equilibrated with the lysis buffer and washed with 10 column volumes of the lysis buffer. Proteins were eluted with the elution buffer (50 mM Tris pH 7.5, 500 mM NaCl, 200 mM imidazole). The eluate was further applied to a 1 mL MBPTrap HP column (Cytiva), washed with 5 column volumes of 20 mM HEPES pH 7.5, 150 mM NaCl, and eluted with 2.5 mL 20 mM HEPES pH 7.5, 150 mM NaCl, 10 mM maltose. The concentration of MBP-hCTD was determined $A_{280}$ using the extinction coefficient $\varepsilon^{MBP\text{-}hCTD}_{280}$ = 143830 $M^{-1}cm^{-1}$. 50 μM MBP-hCTD was incubated with Alexa Fluor 488 NHS

Ester (Invitrogen) with a protein:dye ratio of 1:1.2 at pH 7.5, 25 °C for 2 hours. Free dye was removed by PD-10 desalting columns.

*CTD12.* A DNA sequence encoding 12 tandem repeats of the consensus sequence YSPTSPS was cloned into the pET His6 MBP TEV LIC cloning vector (1M, Addgene plasmid #29656) using ligation independent cloning. This vector encodes a N-terminal 6xHis-tagged MBP and a TEV cleavage site. After TEV cleavage, a serine residue remains at the N-terminus of CTD12, resulting in the protein sequence $S(YSPTSPS)_{12}$, free of cloning artifacts that would otherwise disrupt the consensus sequence. BL21 (DE3) cells containing the CTD12 plasmid were grown in M9 minimal media with $^{13}C$ and $^{15}N$ enrichment achieved through the incorporation of $^{15}N$-ammonium chloride and $^{13}C$ D-glucose (Cambridge Isotope Laboratories) with 50 µg/mL kanamycin at 37 °C and 200 rpm. At an $OD_{600}$ of 0.7-0.8, expression of was induced with 0.5 mM IPTG at 37 °C and 200 rpm for 3 hours. Cells were pelleted by centrifugation at 5000x g and stored at -80 °C. Cell pellets of CTD12 were resuspended in the lysis buffer (50 mM Tris pH 7.5, 500 mM NaCl, 20 mM imidazole) supplemented with 1x EDTA-free protease inhibitor cocktail (Millipore), 1 mM PMSF, and 10 units of RNAse-free DNAse (NEB). Following sonication on ice, the cell lysate was centrifuged at 18000x g for 45 minutes. The supernatant was passed through Ni-NTA resins (Thermo Fisher Scientific) equilibrated with the lysis buffer and washed with 10 column volumes of the lysis buffer. Proteins were eluted with the elution buffer (50 mM Tris pH 7.5, 500 mM NaCl, 200 mM imidazole). The eluate was incubated with TEV protease (1 mg per 1 L growth) under dialysis against 2 L of 50 mM HEPES pH 7.5, 150 mM NaCl at 4 °C for 16 hours. Dialysates were passed through a second Ni-NTA column, and the flow-through containing CTD12 was collected and concentrated using a Amicon Ultra Centrifugal Filter with a 3 kDa molecular weight cutoff (MilliporeSigma). The concentration of CTD12 was determined by $A_{280}$ using the extinction coefficient $\varepsilon^{CTD12}{}_{280} = 17880\ M^{-1}cm^{-1}$.

**Crowder preparation**

Solutions of polymer and protein crowders were prepared by dissolving their powder form in 50 mM HEPES pH 7.5, 150 mM NaCl with vortexing and centrifugation to a final concentration of 400 g/L, except for lysozyme due to solubility issues. 200 g/L lysozyme powder/buffer mixture was incubated in the 37 °C water bath for 30 min to dissolve. Protein solutions were syringe filtered with 0.2 μm membranes. The manufacturer information of each crowder is as follows: poly(ethylene glycol) (PEG) BioUltra, Molecular Biology, 8,000 (Sigma-Aldrich, CAS: 25322-68-3); Ficoll PM70 (GE Healthcare, UNSPSC Code: 41105330); Dextran, from Leuconostoc mesenteroides (Sigma, CAS: 9004-54-0); OmniPur BSA, fraction V, heat shock isolation (EMD Millipore, CAS: 9048-46-8); lysozyme, egg white, ultra pure grade (VWR, CAS: 12650-88-3).

*E. coli lysate.* A seed culture from a single colony of BL21 (DE3) with OD 1 was used to inoculate 30 L LB culture in a Sartorius Biostat Cplus 30 L vessel. Cells were grown at 37 °C with shaking at 250 rpm for 16 hr. The culture was harvested by centrifugation at 35000 rpm in a Sharples T-1-P Super Centrifuge. Cell pellet was resuspended in 2.5x volume of RO water. Resuspended cells were lysed with microfluidization and centrifuged at 38000x g at 4 °C for 30 min. Lysate was lyophilized at -51 °C and 0.120 mBar in a FreeZone 18 Liter Console Freeze Dryer. The lyophilized lysate from 30 L LB culture weighed 18.92 g. The 200 g/L lysate solution was prepared by dissolving 200 mg lyophilized lysate in 1 mL Milli-Q water. The pH of the reconstituted *E. coli* lysate was 7.5.

**NMR data collection and analysis**

NMR data were collected on a Bruker Avance NEO 600 MHz spectrometer equipped with a TCI triple-resonance cryoprobe. All data collection and initial processing were carried out in TopSpin (Bruker). $^{1}$H, $^{13}$C, and $^{15}$N chemical shifts were directly and indirectly referenced to sodium trimethylsilylpropanesulfonate (DSS). All NMR experiments, unless specified, were carried out using 425 μM CTD12 in 50 mM HEPES pH 7.5, 150 mM NaCl, 1 mM DSS, and 10% $D_2O$, with or without 100 g/L of crowder.

$^{13}$C direct detect CON experiments were carried out with 8 scans, 1024 ($^{13}$C) × 256 ($^{15}$N) complex data points, and sweep widths of 10 ppm and 42 ppm for $^{13}$C and $^{15}$N, respectively. Peak picking and assignments were carried out manually using NMRFAM-Sparky. Average chemical shift perturbations were weighted by amino acid types according to the chemical shift standard deviation from Biological Magnetic Resonance Bank as follows:

$$\langle \Delta\delta \rangle = \left\{ \frac{1}{2} \left[ r_{N-C}^{aa} (\Delta\delta_N)^2 + (\Delta\delta_C)^2 \right] \right\}^{1/2},$$

where $r_{N-C}^{aa}$ is the ratio of the standard deviation of the $^{13}C$ chemical shift to that of the $^{15}N$ chemical shift for a specific amino acid type. $r_{N-C}^{Tyr}$ is 0.399, $r_{N-C}^{Ser}$ is 0.875, $r_{N-C}^{Pro}$ is 0.635, and $r_{N-C}^{Thr}$ is 0.191.

**Confocal Imaging**

Glass coverslips (#1.5) were functionalized with a poly(ethylene glycol) silane to increase the contact angle of the droplets for improved visualization[1]. The coverslips were soaked in saturated KOH (Sigma) in isopropanol (Sigma) for 30 minutes, then rinsed thoroughly, and dried overnight at 70°C. 1.2mL 3-[methoxy(polyethyleneoxy)6-9]propyltrimethoxysilane (Gelest Inc.) was diluted to 400mL with anhydrous toluene (Sigma); the dried coverslips were incubated in this solution for 4 hours at room temperature. The coverslips were then washed thoroughly with water and ethanol, and dried overnight at 70°C prior to use.

Confocal images were taken on a Leica TCS SP5 laser scanning confocal inverted microscope with Leica LAS AF software using a HCX PL APO CS ×63.0/1.40 NA oil UV objective. Samples were prepared by mixing 10 µL 56 µM MBP-hCTD, 1 µL 2 µM AF488-MBP-hCTD, 10 µL varying crowder as described above. The final MBP-hCTD concentration was 25 µM with 100 nM AF488-MBP-hCTD. Samples were placed on a functionalized coverslip within a 2.5mm thick imaging spacer, then covered with an unmodified coverslip to prevent evaporation. Samples were allowed to incubate at room temperature on the coverslips for a minimum of 30 minutes prior to imaging to allow the droplets to settle to the glass surface. AF488-MBP-hCTD was excited using the 488nm laser line with a triple dichroic mirror, TD 488/543/633. 2µm polystyrene beads were added to aid in focusing samples that did not form droplets or aggregates.

REFERENCES

(1) Razunguzwa, T. T.; Warrier, M.; Timperman, A. T. ESI-MS compatible permanent coating of glass surfaces using poly(ethylene glycol)-terminated alkoxysilanes for capillary zone

electrophoretic protein separations. *Anal Chem* **2006**, *78* (13), 4326-4333. DOI: 10.1021/ac052121t.
